# Spatial patterns of trematode-induced pits on bivalve skeletons: Challenges and prospects for research on parasite-host dynamics

**DOI:** 10.1101/2025.03.13.643152

**Authors:** Alexis Rojas, John Warren Huntley, Monica Caffara, Daniele Scarponi

## Abstract

Interactions between the parasitic larvae of digenean trematodes (mainly gymnophallids) and bivalves often result in characteristic shell malformations, i.e., pit-like traces. Tracking these traces through the Holocene and modern marine death assemblages has made studying parasite-host responses to natural and anthropogenic environmental change possible. Despite major breakthroughs, empirical explorations of parasite-host dynamics in the geological record are primarily based on trace occurrence data, overlooking that trace spatial patterns on the host skeleton could carry ecological information and potentially document different aspects of the parasite-host interactions (e.g., infective behavior, association with specific host anatomy, spatial relationships of traces with different qualitative properties such as size class, etc.). The Spatial Point Pattern Analysis of Traces (SPPAT) has been increasingly employed to overcome similar challenges in studying predatory traces on bivalve prey. Although this approach holds considerable promise for research on trematode–host dynamics, several assumptions and caveats need to be considered (e.g., the number of traces required to capture the parasite-host dynamics accurately, the reliability of point patterns constructed from multiple host skeletons in describing parasite interactions). Here, we introduce a spatially explicit framework for extracting information from spatial patterns of trematode-induced pits on bivalve shells using SPPAT, address methodological questions involved in assembling a point pattern of traces from multiple host specimens, and discuss critical issues related to drawing inferences from pooled point data. We illustrate our approach using a case study on late Holocene samples of the commercially relevant bivalve *Chamelea gallina* from the northern Adriatic of Italy. This species holds high value in the seafood industry and is increasingly used in climate change research. Our results reveal that trematode-induced malformations on bivalve shells are not random; they show an aggregated pattern for metacercaria traces of the same size classes, while an independent pattern arises when examining traces of two distinct size classes. This study highlights the potential value of spatial information from parasite-induced traces in enhancing our understanding of parasite-host dynamics over time.

## 1 Introduction

Digenean trematodes of aquatic organisms live and reproduce in vertebrate hosts, releasing their eggs through host feces into the water. To complete their life cycle, they must first infect invertebrates as first or second intermediate hosts (Esch, 2002). Digenean trematode infections have major impacts on their hosts, representing a significant population and community structuring driver, particularly in coastal ecosystems (Sousa, 1991; Mouritsen and Poulin, 2002a, 2005). Although different parasites infect marine mollusks, digenean trematodes are the most common metazoan parasitizing these invertebrates in coastal waters (Lauckner 1983; Sindermann, 1990). Studies on parasite-induced malformations on their intermediate macrobenthic hosts indicate that digenean trematodes (typically Gymnophallidae) are primarily responsible for the induction of diagnostic traces in their mineralized skeletons (Ruiz and Lindberg 1989; Huntley 2007). These biotic traces have been commonly preserved in the fossil record since Eocene times (e.g., Benvenuti and Dominici, 1992; Todd and Harper, 2011; Huntley and Scarponi, 2012) but are rarely found in the Late Cretaceous (Rogers et al., 2018). They can potentially document ecological aspects of the trematode–host interactions (i.e., prevalence, diversity, etc.) across changing environments in the space-time continuum. For instance, studying trematode-induced traces in the commercially important bivalve *Chamelea gallina* from both late Holocene and recent shoreface environments of the Adriatic Sea of Italy, before and following significant human impact, uncovered a drastic reduction (by an order of magnitude) in parasite-host interactions, paralleling the rising human influences on the Adriatic and its transition into an urban sea (Fitzgerald et al., 2024). Despite these discoveries, empirical explorations of parasite-host dynamics usually ignore the spatial information inherent to trace location.

Research on spatial patterns in the distribution of parasite-induced malformations in host skeletons remains limited to qualitative assessments of both trace locations and experimental observations. Quantifying this aspect in antagonistic interactions could provide valuable insights into the dynamics between parasites and hosts and help us understand whether their relationships have remained stable through time. Researchers are increasingly employing spatially explicit methods to overcome this common challenge in the study of antagonistic interactions in both fossil and modern records. The location of biotic traces on shelled invertebrates enables spatially explicit analyses (e.g., Rojas et al., 2017; Rojas et al., 2020; Karapunar et al., 2023). For example, research focusing on modern ecosystems (Dietl and Alexander, 2000; Chiba and Sato, 2012) and studies utilizing the fossil record (Kelley, 1988; Dietl et al., 2001) indicate that drilling gastropods targeting bivalve prey often exhibit significant spatial stereotypy. This behavior mirrors the methods of prey handling during the attack and correlates with the morphology of the prey (Kingsley-Smith et al., 2003; Rojas et al., 2015). Approaches used to assess those spatial patterns have varied widely, including qualitative descriptions (Negus, 1975) and a multitude of quantitative methods (Kelley, 1988; Kowalewski, 1990; Anderson et al., 1991; Dietl and Alexander, 2000; Hoffmeister and Kowalewski, 2001; Hammer and Harper, 2024). These methodological inconsistencies make it difficult to compare results across studies in the growing body of research evaluating patterns in the distribution of biotic traces on shelled invertebrates. In addition, most of the methods for analyzing site selectivity in drilling predation were primarily developed to test the null hypothesis of a random distribution of drill holes across arbitrarily defined sectors of the prey skeleton using goodness-of-fit, chi-square or Kolmogorov-Smirnov statistics (Kelley, 1988; Kowalewski, 1990; Anderson et al., 1991), and diversity metrics such as the Shannon-Weaver index (Dietl et al., 2001). These approaches fail to exploit the high-resolution information in the spatial relationship between drill hole locations and depend critically on an arbitrary grid system to split the skeleton into sectors (Kowalewski, 2004; Rojas et al., 2020).

The spatial point pattern analysis of traces (SPPAT) was recently introduced to visualize and quantify the distribution of drill holes on bivalve prey (Rojas et al., 2015; Rojas et al., 2020). This approach includes a morphometric-based collection of spatial information on trace location, kernel density, hotspot mapping of spatial trends, and distance-based statistics for hypothesis testing. Although this approach holds considerable promise for research on trematode–host dynamics, as it allows for standardized investigation of a wide range of ecologic and taphonomic data, several assumptions and caveats need to be considered, including the number of trematode-induced traces required to adequately capture the dynamics of a given parasite-host system, as well as some conceptual and methodological implications of evaluating bivariate interactions in point patterns assembled from multiple specimens (i.e., two sets of different trematode-induced pits jointly considered from multiple individuals of a given host). Here, we evaluate these issues by exploring trematode-induced pits on specimens of the bivalve *Chamelea gallina* found in abundance in cored deposits (i.e., core 240-S8 see Huntley and Scarponi, 2021), that accumulated in the shoreface setting of the late Holocene of the northern Adriatic Sea. Our results highlight the potential of distance-based statistics, univariate and bivariate analyses, and marked analyses for future research on parasite-host dynamics. This study indicates that trematode-induced malformations on bivalve shells do not occur randomly. Instead, they exhibit an aggregated pattern for traces of the same size while showing an independent pattern when considering traces of two different size classes.

## 2 Materials and methods

### 2.1 Chamelea gallina (Linnaeus, 1758) as a model for environmental and parasitic research

*Chamella gallina*, also called the striped venus, is a filter-feeding bivalve widely distributed from the Gulf of Cadiz (Atlantic) to the Black Sea, inhabiting sandy bottoms at depths ranging from 0 to 20 m (Pérès and Picard, 1964; Delgado et al., 2023). The striped venus prefers high-energy seabed and is sensitive to hypoxic and anoxic conditions (e.g., Matozzo et al., 2005). The Adriatic Sea is a key area for this bivalve, as clam harvesting primarily takes place there, and the two primary shellfish resources are the indigenous *C. gallina* and the alien *Ruditapes philippinarum*, introduced from the Indo-Pacific region in 1983 (Breber, 1985). While these two species are economically important in the Mediterranean region, they have been severely impacted by anthropogenic factors, including climate change and antagonistic interactions with invasive species, such as the blue crab (Prado et al., 2024; Cabiddu et al., 2025). Adriatic *C. gallina* has experienced a significant decline in harvesting over the last decades of the previous century, falling from 80,000 to 40,000 tons per year, with current production stabilizing at approximately 20,000 tons yearly (Italian Ministry, 2022). Given its decline in ecological and economic significance, numerous studies have been conducted over the years to establish practical tools and diagnostic features for monitoring the present and forecasting the status of this clam resource. A network-based assessment of bibliographic data (figure 1) reveals that research on *C. gallina* is dominated by a large cluster focusing on major relevant study areas for *C. gallina*. This research landscape is further complemented with highly interconnected minor clusters dedicated to food safety and processing, current threats, and physiology related to changing environments and associated with the other shellfish resources (i.e., *Ruditapes philippinarum* figure 1). This highlights the significant value of *C. gallina* in the seafood industry and marine environmental research. The network analysis of literature data (figure 1) highlights the general focus on modern systems, suggesting a gap in research on detecting long-term ecological responses to past climate-driven environmental changes that extend beyond the limited timeframe of direct ecological monitoring. In this respect, for this species few studies have attempted to bridge the paleontological-ecological gap concerning anthropogenic and climate-driven environmental changes, with notable exceptions including Huntley and Scarponi (2015) on *C. gallina* parasitism, Cheli et al. (2021) on shell microstructure, and Scarponi et al. (2023) on community changes in *C. gallina* associations before and during the Anthropocene (*sensu* Crutzen, 2002).

**Figure. 1.**
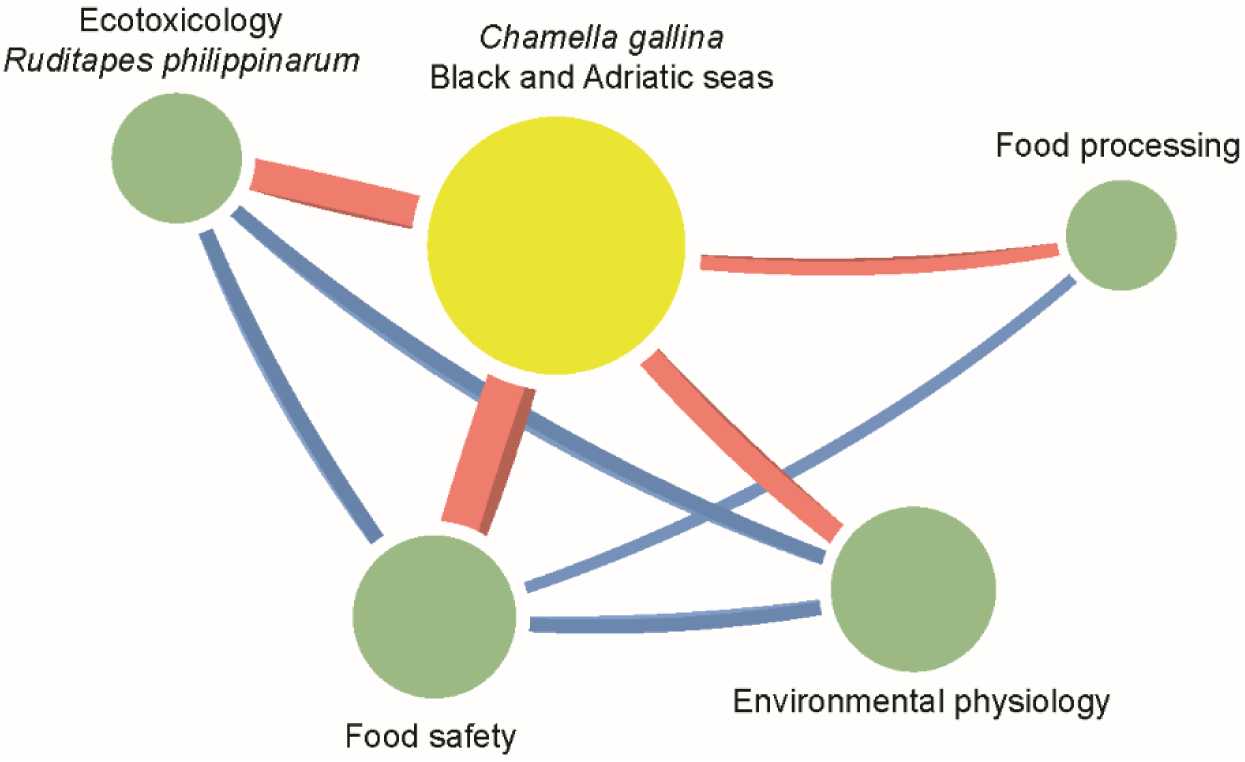
Research landscape of *C. gallina*. The figure shows the modular structure of a co-occurrence network, including 49 keywords connected through 265 links (Table S1). Bibliographic data was compiled from 349 documents in the Web of Science using the search query: (Chamelea OR Venus) AND (gallina). The network was partitioned into modules using the Map Equation framework, with each module interpreted as a research area inferred from the keywords. Modules are represented as circles, with areas proportional to their size (i.e., number of keywords) and aggregated inter-module links with widths proportional to their connections. The yellow circle highlights the research area where the keyword “Chamella gallina” is clustered. The analysis and visualization were created using the Map Equation framework described in Rojas et al. (2024).

In the *C. gallina* parasite-host system context, the striped venus serves as a second intermediate host for several species of digenean trematodes. Bartoli (1974) and Bartoli and Gibson (2007 and references therein) recognized at least two gymnophallid species parasitizing *C. gallina* as second intermediate host*: Gymnophallus rostratus* and *Parvatrema duboisi*. Another trematode, *Himasthla quissetensis*, which affects *C. gallina* as a metacercaria and is part of the Echinostomidae family, has also been documented (Bartoli and Gibson, 2007). However, *Himasthla* cercariae penetrate the gills of their second hosts and, in a few hours, transform into metacercariae by forming a thin-walled cyst in the gills or foot of various bivalves including also *C. gallina* (Stunkard, 1938; Belousova, 2023). However, there have not been any documented traces on the inner surface of the bivalve connected to this encystment of *Himasthla* metacercariae. Indeed, there is a general dearth of information on the morphology of host shell growth response to trematode parasites in the neoparasitological literature, so, in many cases, we must make assumptions based on what we know from closely related gymnophallid taxa. *Gymnophallus rostratus* has a complex life cycle involving multiple potential host taxa. *Loripes lacteus*, a bivalve (Bartoli 1974, 1982), has been documented as *the first intermediate host* of *G. rostratus*. From there, the parasite undergoes asexual reproduction, multiplying into numerous cercariae, the free-swimming larval stage. These cercariae are then released into the water column, where they actively seek out their second intermediate host – including *C. gallina* and nearly a dozen other bivalve genera (Bartoli, 1974). Within these bivalves, the cercariae reach the extrapallial space within the pallial line, transforming into metacercariae, which, as in many gymnophallids, feed nutrient-rich fluids and tissues from the host waiting to fully develop in the avian definitive host’s digestive system (i.e., *Aythia ferina*; Bartoli and Gibson, 2007). However, with respect to other gymnophallid infecting *C. gallina*, this species does not seem to elicit any pallial reaction towards the metacercariae, even when these are very numerous (Bartoli, 1982). Once inside the bird’s digestive system, the metacercariae excyst and mature into adult worms, primarily in the bird’s intestine. Here, they reproduce sexually, releasing eggs that are shed back into the environment through the bird’s feces, restarting the cycle.

Similarly, *P. duboisi* relies on multiple hosts during its complex life cycle. Bartoli (1974) could not identify the first intermediate host but confirmed the presence of *P. duboisi* metacercariae in *C. gallina*, *M. galloprovincialis*, and *Brachydontes minimus*. Only recently *Ruditapes philippinarum* has been recognized to be the first and second intermediate host of *P. duboisi* in Korea (Jung et al., 2021). Concerning the definitive hosts, *Aythya fuligula* (Bartoli, 1974) and recently *Calidris tenuirostris* (Jung et al., 2021) were confirmed as natural definitive host of *P. dubosi*; Bartoli was able to infect *Gallus gallus*, *Anas platyrhynchos*, and *Larus argentatus michaellis* with the same gymnophallid. Montenegro et al. (2021) reported the presence of concave holes with orange-brownish coloration in the internal parts of the shell of *Leukoma theca* due to unidentified *Parvatrema* metacercariae. In this regard, the literature emphasizes the necessity of further explore the antagonistic interactions of *C. gallina*, as there is limited understanding of trematode parasites’ activities in living *C. gallina* and their host responses.

### 2.2 Spatial Point Pattern Analysis of Traces (SPPAT)

#### 2.2.1 Extracting point data on parasitic traces

##### Landmark detection

The spatially explicit analysis of parasitic traces requires assembling a dataset of their locations on the host skeleton. Following previous research on predatory traces on bivalve prey, we employed a two-dimensional morphometric approach to quantify trace position (Roopnarine and Beussink 1999). Images of the specimens, in internal view and oriented perpendicular to the commissural plane, were obtained with a digital camera Nikon D5300 and AF-S DX Micro-NIKKOR 40mm f/2.8G Close-up Lens, attached to a Kaiser Copy Stand System. All images were oriented using Adobe Photoshop 2024 such that the anteroposterior axis, proxied by the ventral limit of the scars formed by the adductor muscles (Figure 2), was horizontal to facilitate consistency in landmark data collection (Kolbe et al., 2011). Images of the right valves were flipped horizontally to enable the use of all shell material in the analysis.

**Figure 2.**
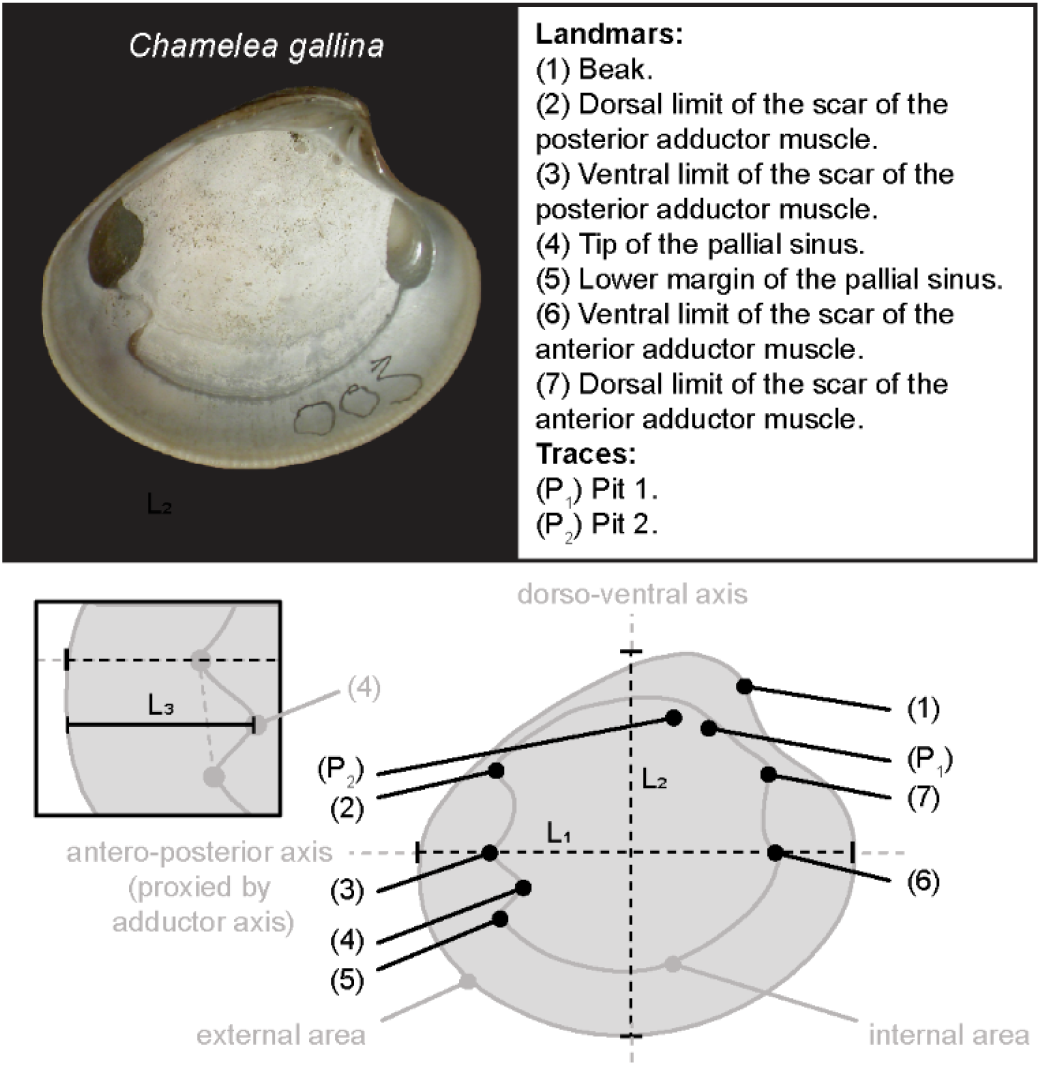
Sample image and corresponding landmarks for point data detection and linear measurements taken on C. gallina specimens. L1: valve length; L2: valve height; L3: pallial sinus index.

Seven landmarks were selected to capture the shape of the host, six of them following a previous study documenting differences in shape between *C. gallina* and *C. striatula* from the Portuguese coast and conducted to resolve taxonomic issues (Rufino et al., 2006) (landmarks 1 to 6). An additional landmark representing the upper limit of the anterior adductor muscle scar was also selected (landmark 7) following a recent study of point-based homologies across a range of bivalve morphological groups (Edie et al., 2022).

Because trematode-induced malformations can occur on either the left or right valves of the hosts, we excluded landmarks located along the hinge, as they are not recognizable on both sides. This choice represents a compromise between accurately describing the host shape and maximizing the parasitic traces obtained from the fossil samples. The center of the pits, defined by the intersection of its major and minor axes (*sensu* Fitzgerald et al., 2024), was used to register trace locations and treated as pseudo-landmarks. Point data, i.e., *x* and *y* coordinates of pixels on the images, corresponding to the landmarks and traces, were manually detected using the *locator* R-function.

##### Geometric morphometric analyses

Landmarks and pseudo landmarks positions were rotated, scaled, and translated through the Bookstein baseline registration method for two-dimensional data (Bookstein 1991) using landmarks 3 and 6 located on the anteroposterior axis (Rojas et al., 2020). The R functions for landmark-based morphometrics of Claude (2008) were used to carry out this analysis. The dataset is presented in the Supplementary Data S1.

#### 2.2.2 Building a point pattern dataset of parasitic traces

##### Observation window

The point pattern analysis of trematode-induced pits requires designating the region where the parasitic traces are recorded. Following the approach for landmark detection described above, we digitalized the valve perimeter (in the form of a series of data points) of a well-preserved adult specimen and named this region outer area (Figure 2). We added two additional points to this outer area, which correspond to the two landmarks defining the baseline used for registration (landmarks 3 and 6) (Edie et al., 2022), required to overlap the traces and the observation window to create a point pattern. In addition, we digitalized the internal area of the valve, defined as the region between the muscle scars, limited ventrally by the pallial line and dorsally by the hinge area. These two areas were selected because they help describe spatial patterns in the distribution of the traces and can be unambiguously identified in the photographs. The outer area was used as an observation window for the point pattern, with the inner area outlined as a reference. The dataset of *x* and *y* coordinates of vertices of the external area was rotated, scaled, and translated using the two additional points as a baseline. The outer area is presented in the Supplementary material 2.

##### Marks

Point patterns can have marks or additional attributes attached to each point (Baddeley et al., 2016). We use linear measurements describing the size of the trematode-induced traces examined to create the *Pit size* mark defined as the geometric mean of the major and minor axes of the pits. It is used to create a categorical mark with the levels “small” and “large”. The methodological decision to create two size classes, as well as the particular threshold (0.55 mm), follows a previous study of the modern and late Holocene samples included in our research, indicating that both sets of pit samples are made up of two Gaussian distributions (Fitzgerald et al., 2024).

##### Planar point pattern

We constructed a marked point pattern by combining the coordinates of the parasitic traces, the observation window representing a standardized host skeleton (i.e., study area in Bookstein shape units), and the categorical (host body size and pit size) marks attached to each trace point. In practice, we created an object of class “ppp” (planar point pattern) using the *spatstat* R package version 3.3-0 (Baddeley et al., 2016).

#### 2.2.3 Hypothesis testing

Null models and point process models are tools used in spatial point pattern analysis to examine ecological hypotheses (Velázquez et al., 2016). The choice of a null model in spatial point pattern analysis of biotic traces depends on the underlying research question regarding how the pattern of traces was generated. Complete spatial randomness (CSR) is the simplest theoretical model suitable for evaluating the spatial distribution of biotic traces on shelled invertebrates, including drill holes and parasite-induced malformations. However, different theoretical null models are appropriate for point patterns of trematode-induced malformations on bivalve hosts, depending on whether the parasitic traces belong to the same category (e.g., parasite size class) (Figure 3).

**Figure 3.**
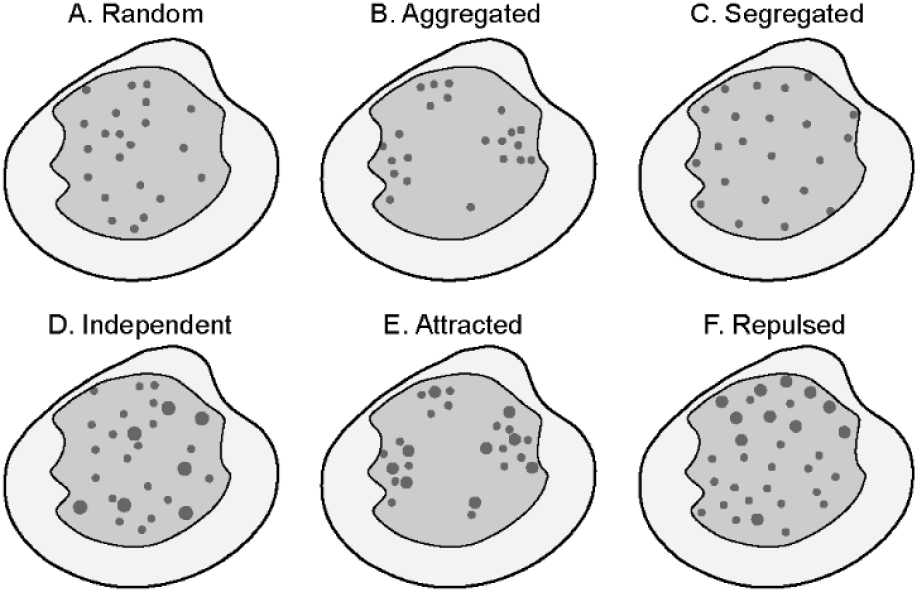
A diagram of idealized point patterns of trematode-induced pits on the skeleton of Chamelea gallina. (A-C) The univariate case considers a set of traces. (D-E) The bivariate case considers two sets of traces with different properties or marks (e.g., parasitic traces of small and large size classes).

In the univariate analysis, which considers a single set of traces, CSR is the appropriate model for analyzing the spatial distribution of trematode-induced malformations and drill holes in bivalve prey (Rojas et al., 2015; Karapunar et al., 2023). This theoretical model can be shown as a homogeneous Poisson process, which assumes that the trace intensity λ is constant across the internal surface of the bivalve skeleton (Figure 3A). Here, we assume the internal shell surface lacks heterogeneity, and therefore, any non-homogeneity in the distribution of traces revealed by distance-based statistics suggests underlying interactions between point events. However, non-homogeneity in the distribution of traces could also point to anatomical features of the bivalve or specific processes controlling the spatial distribution of traces. A point pattern can be described at a given scale as aggregated or segregated (Figure B-C; Baddeley et al., 2016). In bivariate analysis, which considers two sets of biotic traces, the independence null model (Figure 3D) is suitable for investigating the spatial relationship between traces of different trematode size classes (or taxa). This theoretical null model assumes that the two spatial point patterns of traces are generated by independent processes, implying that points of different types do not interact (Wiegand and Moloney, 2004). However, if distance-based statistics reveal a spatial dependence between points of various types, this dependence may result from either attraction (Figure 3E), where points are more closely clustered than expected under independence, or repulsion, where points are more dispersed (Figure 3F; Ben-Said, 2021). For all analyses, the empirical curves of distance-based statistical functions are compared to the Monte Carlo envelopes generated through simulations of the null model (Wiegand and Moloney, 2004). A departure from the null model is indicated when the empirical curves fall outside the simulation envelopes (Badeley et al., 2016). To test for the significance of this departure, we applied a goodness-of-fit (GoF) test (Loosmore and Ford, 2006).

## 3 Results

### 3.1 Reliability of trace point patterns assembled from multiple specimens

Following a baseline registration approach (Bookstein, 1991), we created spatial point patterns of parasitic traces using an undeformed observation window representing the internal surface of the bivalve host. Alternative two-point superpositions of the trace data and observation window (Figure 4) were generated using different pairs of landmarks (Figure 2). This approach allows us to visually evaluate the influence of the baseline selection on the spatial point patterns derived from trematode-induced pits recorded in different specimens of *C. gallina*. Specifically, we used traces registered in four hosts, one per specimen, and obtained eight spatial point patterns in Bookstein-shape coordinates.

**Figure 4.**
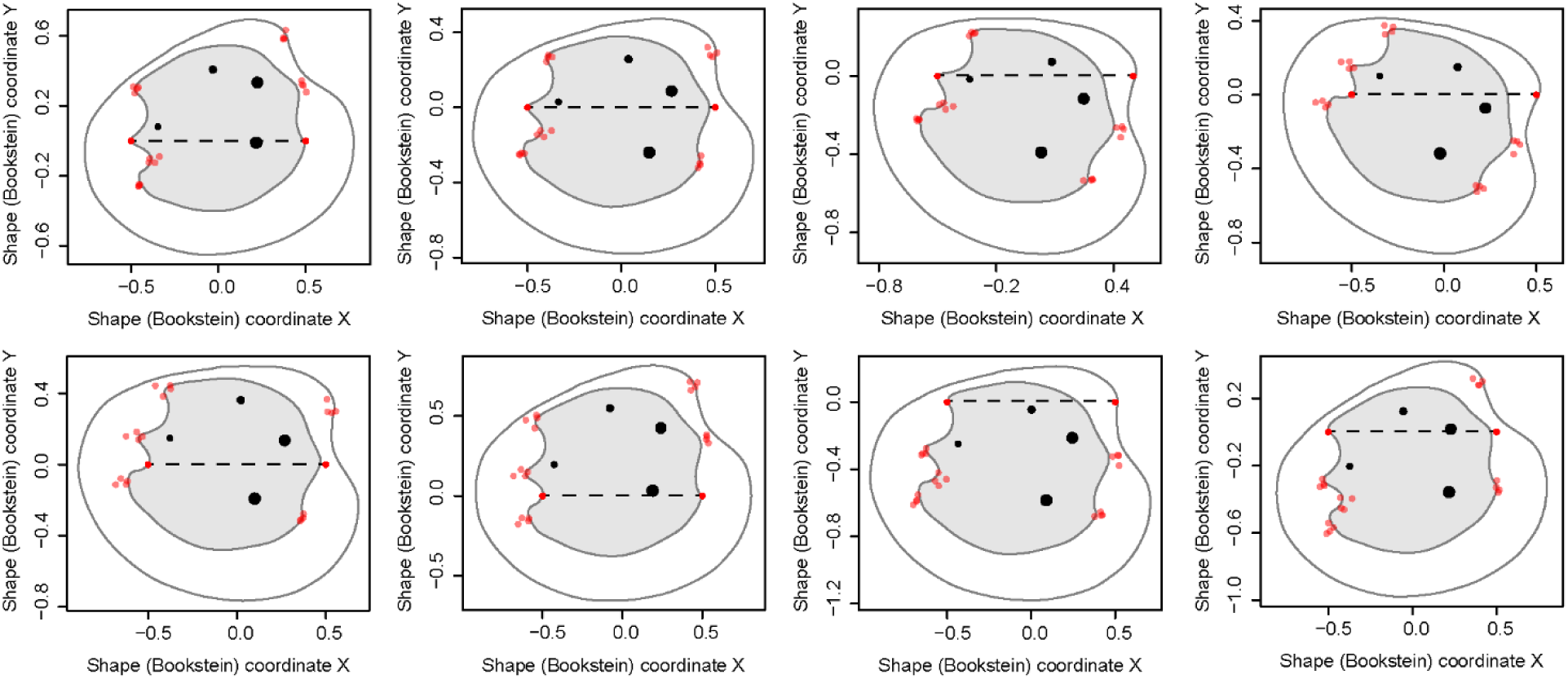
Impact of the baseline selection on the obtained point pattern of trematode-induced pits on the skeleton of C. gallina. Specimen numbers: 13.1.160, 13.1.237, 13.1.310, and 13.1.38.

### 3.2 Point patterns of traces recorded in a single host specimen

SPPAT was applied to parasitic traces recorded within a single host specimen. Distance-based statistics designed for univariate analysis were employed for traces belonging to a single type, such as a specific trematode size class. When traces were categorized into two distinct size classes, bivariate statistical methods were used to analyze the spatial relationship between the two types.

#### 3.2.1 Univariate analysis

Since all the multi-parasitized valves (i.e., those valves with more than one trace) examined in our study exhibit both small and large pits, this univariate analysis was conducted on the two specimens in which most of the traces belong to the same size class: valves 193 R at 13.10m and valve 197 L at 13.10m from core 240S8 (Figure 5). The chosen valves mainly show small pits (18 and 22 traces) but also contain one and four large traces, respectively, all categorized within the small class type for the univariate analysis. Kernel density maps show that the density of traces varies over the shell surface, with the higher density located towards the anterior and dorsal regions of the shell. Because distances are expressed in Shape Bookstein units, derived from the baseline registration analysis, the interpretation of the radio (*r*) in the graphical output of the distance-based statistics should be made in reference to the Kernel Density maps. Although running the SPPAT for multiple traces recorded in a single shell does not require morphometrics (see Figure S1), we conducted baseline registration to ensure consistent reporting of results and to facilitate comparison with the various graphical outputs presented in later sections. The Diggle-Cressie-Loosmore-Ford (DCLF) test of Complete Spatial Randomness (CSR) was conducted separately for specimens 240S8 13.10m 193 R (u = 0.022323, rank = 1, *p*-value = 0.001) and 240S8 13.10m 197 (u = 0.018622, rank = 1, *p*-value = 0.001).

**Figure 5.**
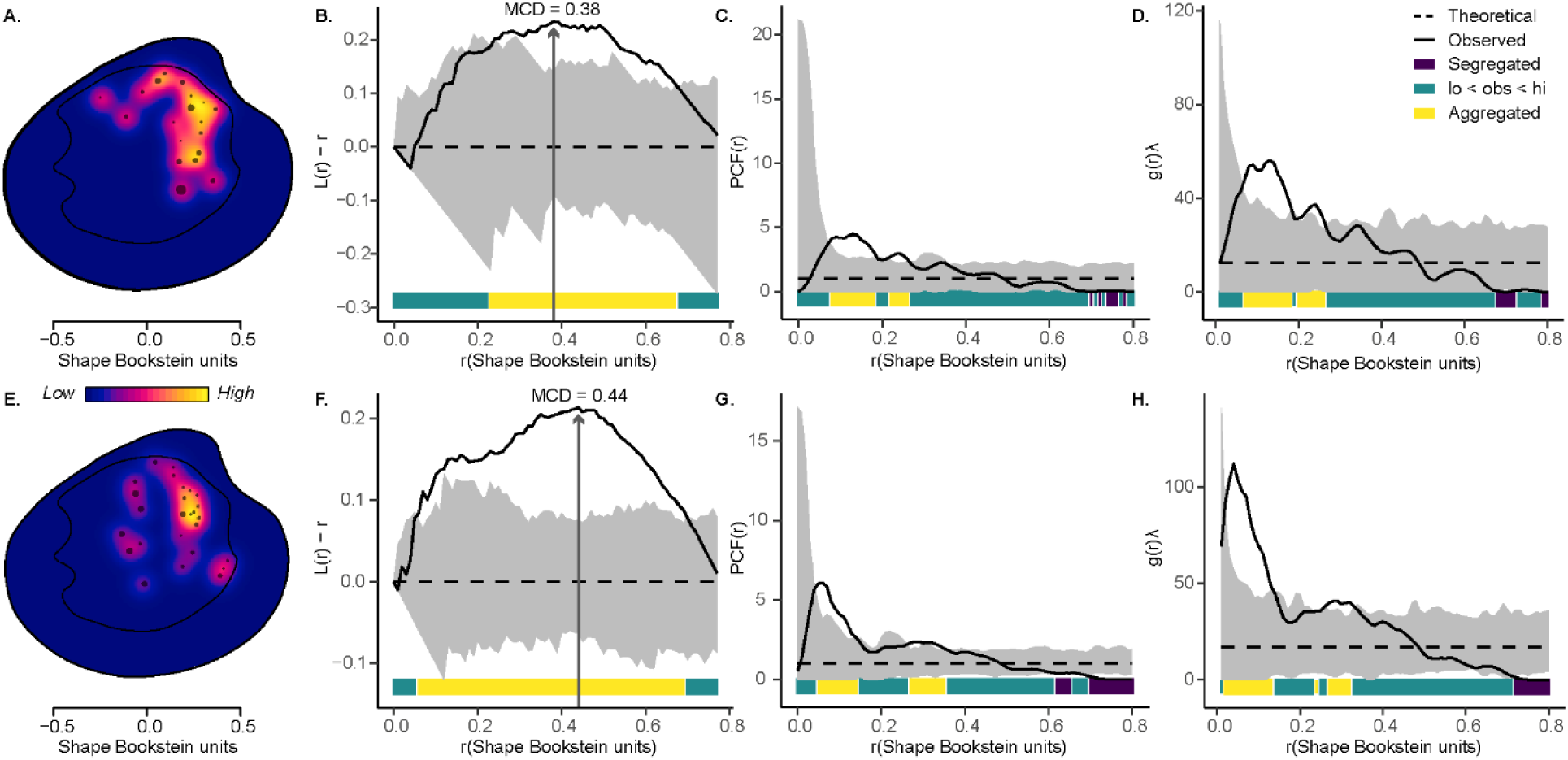
Univariate analysis. The assembled datasets give the locations of trematode-induced pits in one host valve. A-D: Valve 240S8 13.10m 193 R; 19 pits. A. Point pattern and kernel density map. B. L-function. The black arrow indicates the point of maximum clustering distance (MCD). C. Pair correlation function (PCF). D. O-ring function. E-H: Valve 240S8 13.10m 197 L; 26 pits. E. Point pattern and kernel density map. F. L-function. G. Pair correlation function (PCF). H. O-ring function. The dark gray area is the simulation envelope for 999 Monte Carlo simulations of Complete Spatial Randomness CSR (Theoretical). The empirical curves of the distance-based statistical functions (i.e., observed data) are compared to the Monte Carlo envelopes generated through simulations of the null model. A departure from the null model is indicated when the empirical curves fall outside the simulation envelopes. The pattern is aggregated when the empirical curve falls above the envelope; when it falls below, it is segregated. The maximum clustering distance (MCD) represents the scale at which pits are most strongly clustered (i.e., the distance r at which the observed value of the L-function deviates the most from the expected value under CSR). The position of the empirical curve in relation to the simulation envelope of the null model is indicated by a color bar at the bottom. The distance-based spatial statistics were computed using the spatstat package in R (Baddeley et al., 2016), and their graphical outputs were created using the plot quantums function from the ggplot2 package (Wickham, 2016).

The results indicate that the spatial distribution of small and large trematode-induced pits on the examined shells does not follow a random pattern (Figure 5 A-E). This is confirmed by the graphical output of the distance-based *L*-function indicating significant aggregation of pits at short distances, between approximately 0.2 and 0.6 Shape Bookstein units, reaching the maximum clustering (MCD) at a distance of 0.38 and 0.44 in each case (Figure 5 B-F). Looking at the density maps, an MCD equal to 0.4 units represents in the standardized shell approximately half of either the length or height of the internal area (Figure 5 A, E). Furthermore, the pair correlation function (PCF) and the *O*-ring function suggest marginally significant segregation of pits at large distances (> 0.6 Shape Bookstein units) (Figure 5 C-D, G-H) due to grouped distributions where pits form clusters arranged in a linear fashion (Figure 5A, E).

#### 3.2.2 Bivariate analysis

This bivariate analysis examines a spatial pattern with large and small trematode-induced malformations recorded on a single multi-parasitized shell. For this analysis, we selected the valve with the highest number of traces from both size classes (valve 332 R from core 240S8 at 13.10 m) (Figure 6A-B). This experiment tests the null hypothesis of spatial independence between small and large traces (e.g., their locations are not influenced by each other) via random relabeling and using the cross-correlation function (*Kcross*). Specifically, we performed a random permutation of the “small” and “large” traces while keeping both the spatial locations of the traces and their proportions fixed. The graphical output of the distance-based *Kcross* function indicates that the distribution of the two sets of traces on the selected specimen is consistent with the null hypothesis of random labeling, and there is no evidence of spatial distinction between them (Figure 6C-D). These findings are further supported by the segregation test (T = 0.39267, p-value = 0.211), which confirms the lack of significant spatial interaction between small and large pits.

**Figure 6.**
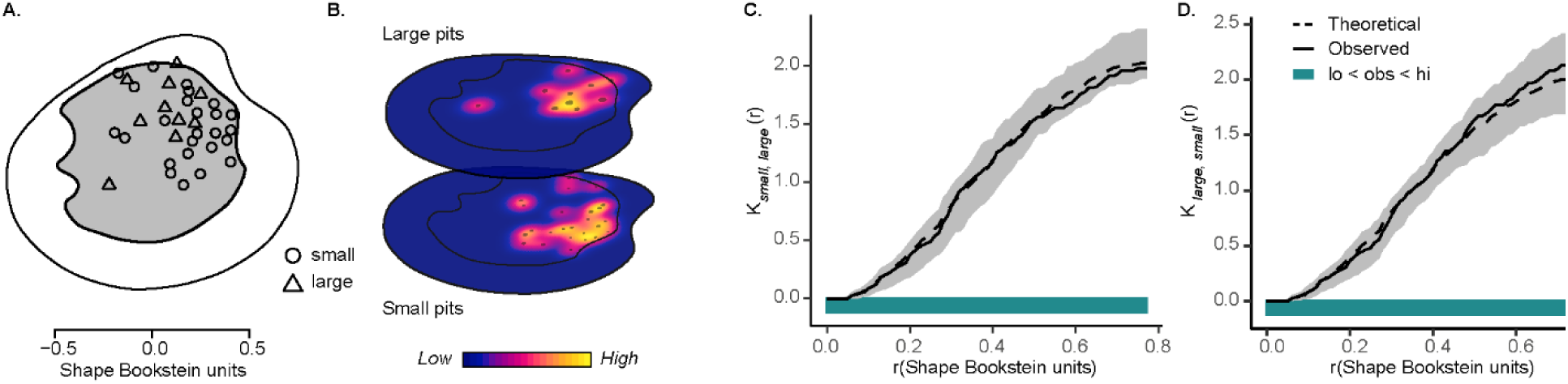
Bivariate analysis. The dataset gives the locations and size of trematode-induced pits in one host valve (valve 332 R from core 240S8 at 13.10m core depth showing 25 small and 10 large pits). Trace size is a categorical variable with two levels. A. Marked point pattern with 35 trematode-induced pits subdivided by their size. B. Kernel density maps for pits of each size category. C-D. Cross-correlation function. The dark gray area is the simulation envelope for 999 Monte Carlo simulations under the null hypothesis of spatial independence between small and large traces (i.e., their locations are not influenced by each other) via random relabeling.

### 3.3 Point patterns combining traces from multiple host specimens

Our experiments examining traces occurring in a single skeleton resulted in a consistent pattern showing significant aggregation at smaller scales, which become segregated, but only marginally significant, at larger distances. Despite this result, such an approach limits the number of traces available for quantitative analysis, raising the question of whether similar trends would emerge in point patterns derived from pooled trace data across multiple skeletons. To address this challenge, we created larger point patterns by combining traces from multiple host specimens. First, we independently performed a univariate analysis of the pattern obtained from combining trace data from all single-parasitized shells (i.e., all skeletons with one trematode trace) with large and small pits. Then, we performed a univariate analysis of the patterns obtained by assembling traces (independently of their size) from all valves bearing the same number of traces, starting from combining valves with one trace up to valves with 63 traces. The latter is the maximum number of traces recorded in a single shell in the studied material. The specific combination of specimens used to create a point pattern depends on the research hypothesis being tested.

#### 3.3.1 Univariate analysis of pooled data from valves with a single pit

Kernel density maps show that higher densities of both large and small traces are concentrated near the dorsal region of the shell (Figure 7). Consistent with our previous experiments, the DCLF test of Complete Spatial Randomness (CSR), conducted separately for the point patterns of small (u = 0.015182, rank = 1, *p*-value = 0.001) and large pits (u = 0.036416, rank = 1, *p*-value = 0.001), provides strong evidence that these patterns deviate from CSR. Regardless of their size, distance-based functions indicate that trematode-induced malformations on *C. gallina* are significantly aggregated at short distances and exhibit marginally significant segregation at larger distances. The segregated pattern observed at larger scales is due to the distinct clusters of pits arranged linearly (Figures 7A, E).

**Figure 7.**
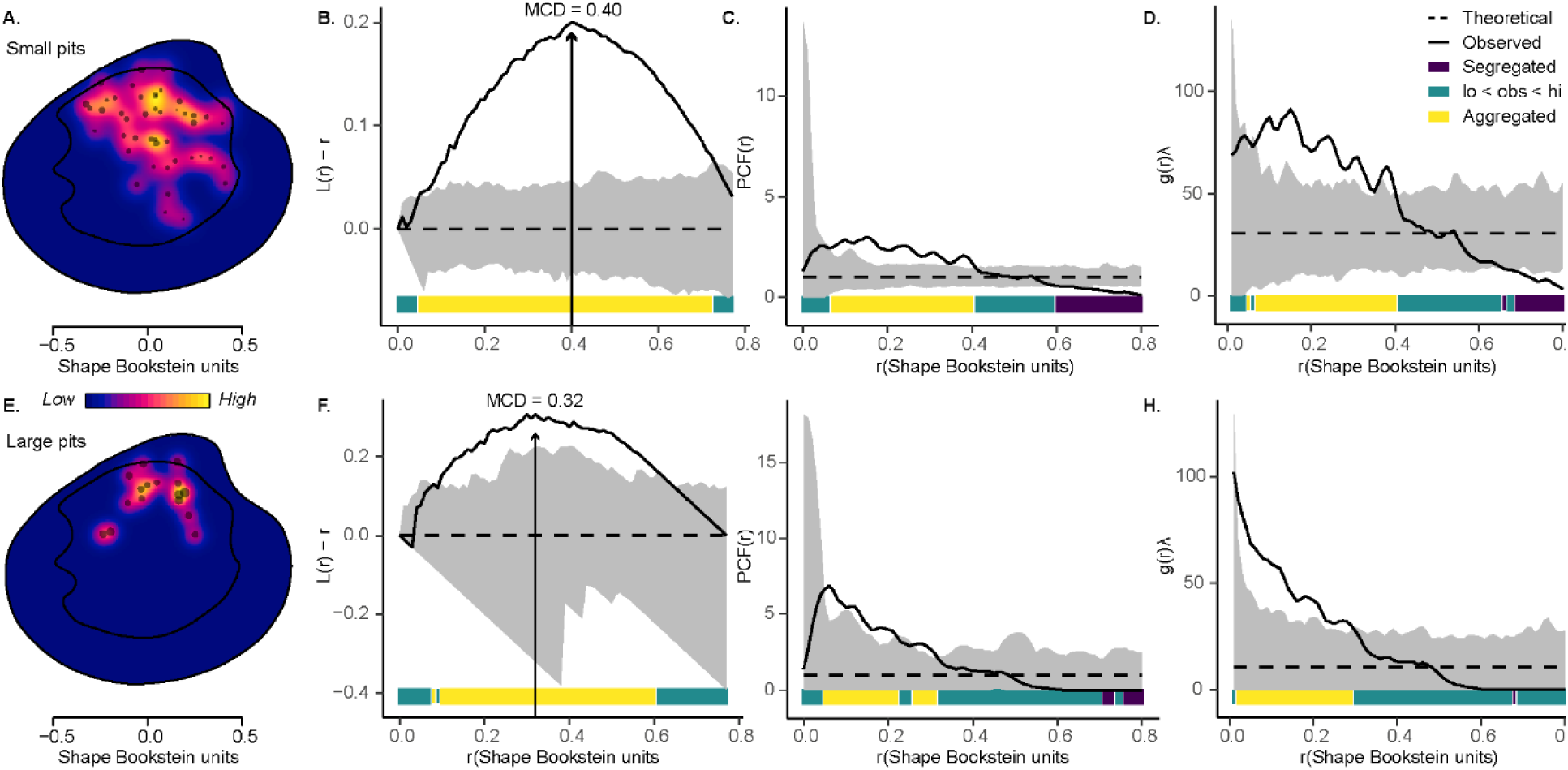
Pooled data from individual hosts with a single trace (single-parasitized shells). The assembled datasets give the locations of trematode-induced pits with the same qualitative property from all hosts exhibiting a unique pit (core 240S8). A-D: All valves with a single small pit; 56 valves. A. Point pattern and Kernel density map. B. L-function. The black arrow indicates the point of maximum clustering distance (MCD). C. Pair correlation function (PCF). D. O-ring function. E-H: All valves with a single large pit; 17 valves. E. Point pattern and Kernel density map. F. L-function. The black arrow indicates the point of maximum clustering distance (MCD). G. Pair correlation function (PCF). H. O-ring function. The dark gray area is the simulation envelope for 999 Monte Carlo simulations of CSR.

#### 3.3.2 Univariate analysis of pooled data from valves with multiple pits

This experiment investigates whether a particular number of traces on a given host influences the spatial pattern while also generating patterns with varying numbers of traces. Given that bivariate patterns of small and large pits in our previous experiments on single multi-parasitized shells (Section 3.2.2) were consistent with the null hypothesis of random labeling, and no evidence of spatial distinction between size groups was observed (Figure 3D), bivariate analysis is not performed on point patterns combining data from multiple specimens. This observation also supports our decision to conduct univariate analysis on point patterns that combine both size groups. Specifically, we conducted a univariate analysis of 23-point patterns from specimens in the entire dataset by grouping trace data from all valves bearing the same number of traces. This analysis starts with valves exhibiting a single trace (single-parasitized shells analyzed in Section 3.3.1) and continues with those showing 2, 3, 4 traces, and so on. In each case, the DCLF test of complete spatial randomness (CSR) provides strong evidence that they do not follow CSR (Table S1). Although there is a large variation in the number of valves with a given number of traces (e.g., 62 valves exhibit a single pit, whereas only one valve exhibits 63 pits), distance-based functions indicate that trematode-induced malformations follow a similar spatial pattern, showing aggregation of pits at short distances and segregation at large distances (Figure 8). However, the results suggest that the total number of traces in the point pattern, regardless of whether they originate from individuals with single or multiple parasitic pits, ultimately influences the observed spatial pattern. This raises the question of how many traces are necessary to fully capture the dynamics of the studied system. We address this question in the experiment described below.

**Figure 8.**
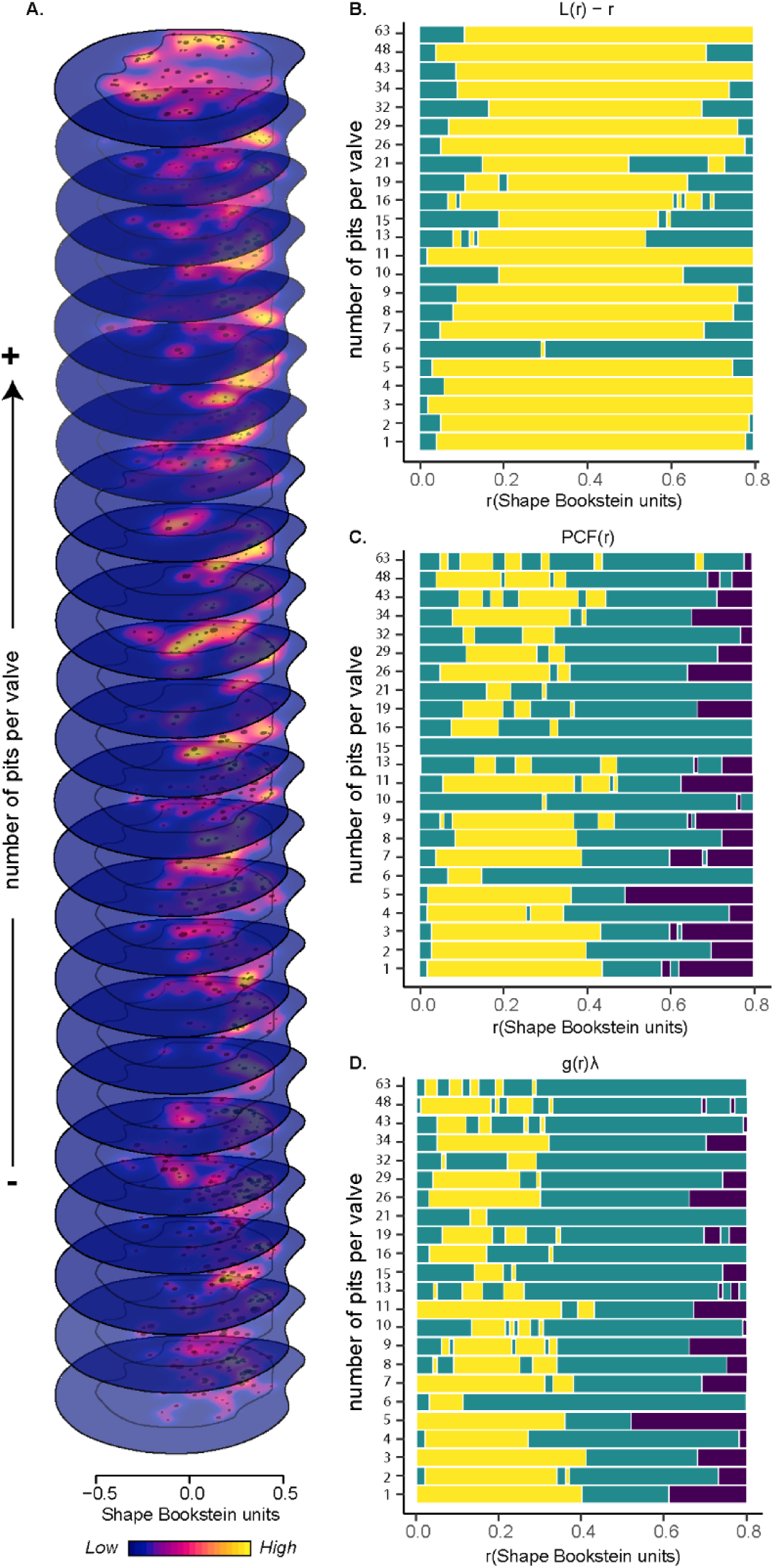
Pooled data from hosts with multiple traces (multi-parasitized shell). The assembled datasets give the locations of the trematode-induced pits from all valves exhibiting the same number of traces (core 240S8). A. Point patterns and Kernel density maps. B-D. Stacked color bars indicate the position of the empirical curves relative to the simulation envelope of the null model in each case. B. L-function. C. Pair correlation function (PCF). D. O-ring function. The simulation envelopes for 999 Monte Carlo simulations of CSR are not shown for simplicity. Distance-based statistics suggest aggregation at medium to short distances and segregation at larger distances (see Table S1).

### 3.4 Bootstrapped Spatial Point Patterns

In this experiment, we sequentially assembled point patterns representing the locations of trematode-induced pits through bootstrap resampling of point data from core 240S8. These patterns ranged from 15 to 100 traces, increasing by five traces per interval. We generated 10 point patterns for each increment and estimated the pair correlation function for each. As a graphical output, we overlaid the 10 color bars that summarize the position of the empirical curves relative to the simulation envelope of the null model for each bootstrapped point pattern (Figure 9). Similar to our previous experiments, the results indicate that the pattern describing trematode-induced malformations *C. gallina* is significantly aggregated at short to intermediate distances, with pits clustered around the dorsal region of the skeleton. However, at larger distances, the pattern transitions to significant segregation, resulting from distinct clusters of pits that are spatially separated (Figure 9B). This significant segregation of traces begins to emerge in some bootstrapped point patterns when approximately 20 pits are considered (see also Figure S1) but becomes consistently present across all bootstrapped patterns once approximately 35 traces are included.

**Figure 9.**
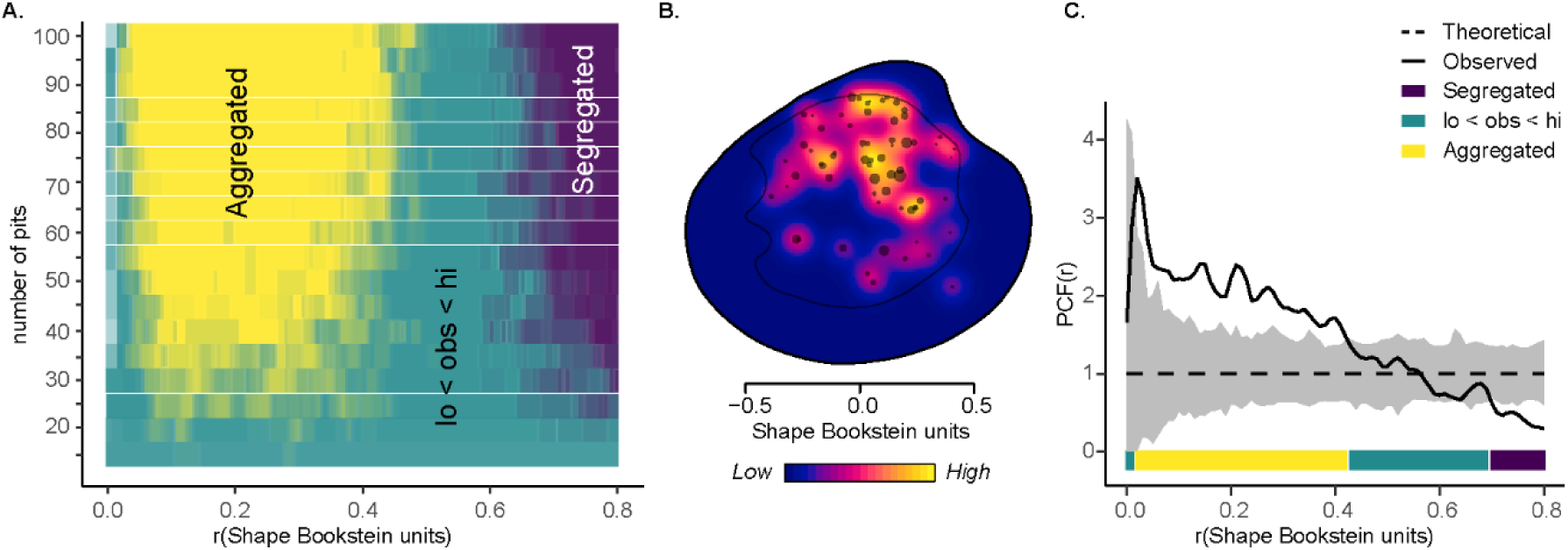
Bootstrapped spatial point patterns. The assembled datasets give the locations of the trematode-induced pits obtained through bootstrap resampling of point data from core 240S8, ranging from 15 to 100 traces with increments of 5 traces per interval. A. Bars from the graphical output of the Pair Correlation Function (PCF). Each interval includes 10 overlapped bars (with 20% opacity) representing the same number of bootstrapped point patterns. The simulation envelopes for 999 Monte Carlo simulations of CSR for each bootstrapped point pattern are not shown for simplicity. B. Point pattern and Kernel density map for a resampled pattern with 70 pits. C. Pair correlation function (PCF) estimated for the resampled point pattern in B. The dark gray area is the simulation envelope for 999 Monte Carlo simulations of CSR.

Nevertheless, the same pattern persists across a wide range of trace numbers. Overall, this experiment confirms that the high density of traces located around the dorsal portion of the shell underlies the significant aggregation observed at medium to short spatial scales in the graphic outputs of the distance-based statistics. In addition, the occurrence of multiple clusters of pits, located, for instance, near the ventral edge of the shell in the Kernel density map illustrated in Figure 9B, produce the segregated pattern observed at larger scales (approximately > 0.6 Shape Bookstein units that represent in the standardized shell more than half of either the length or height of the internal area) (Figure 9A). Furthermore, this experiment confirms that such a segregated pattern is statistically significant (Figure 9C) and requires a certain number of point data to be captured (approximately > 20 traces).

## 4. Discussion

### 4.1 Methodological challenges and prospects for research on parasite-host dynamics

The first methodological challenge in the spatial point pattern analysis of parasitic traces on bivalve host is designating the proper observation window (Baddeley et al., 2016). Here we considered the outer area of an inner valve as well as the internal area (as defined in section 2.2.2) because they can both be unambiguously identified in the photographs. Given that parasitic-induced malformations are generally infrequent outside the internal area, it can serve as an observation window. However, trematode-induced malformations have been noted beyond this area, particularly along the ventral edge, siphonal region, and around the umbonal area, which is partially obscured by the hinge in the examined photographs (Figures 7-9). Thus, because all regions of the valve can potentially exhibit a trace, the entire inner surface of the valve (outer perimeter) was used as an observation window for the point pattern. Indeed, despite potential individual variations, performing SPPAT requires the use of a single, fixed polygonal region, that should be identified with the sum of internal and external areas (so the outer perimeter). We employed the outer perimeter of an adult specimen and applied the baseline registration approach (i.e., a reference line from landmarks 3 to 6; Figures 2, 4A). Although Procrustes shape coordinates and thin-plate splines are currently more popular approaches for geometric morphometric analyses (Bookstein, 2023), by applying the baseline registration we obtained an undeformed observation window, i.e., it preserves the relative proportions of the bivalve shell, that can be aligned with the trace data and facilitate the interpretation of the spatial patterns. Alternative two-point superpositions of the trace data and observation window were generated using different pairs of landmarks (Figure 2) to visually evaluate the influence of the baseline selection on the spatial point patterns derived from trematode-induced pits recorded in different specimens of *C. gallina*.

Our experiments show that assembling a spatial point pattern of trematode-induced malformations from multiple traces recorded in a single skeleton (i.e., a multi-parasitized shell) does not require morphometric methods to standardize shape and size, allowing researchers to report the graphical outputs in standard units (e.g., mm or cm) (Figure S1). The main requirement is to employ a specimen with a number of traces that allow computing distance-based functions for hypothesis testing and that fully capture the overall pattern (see Figure 9A). Remarkably, these point patterns already clearly represent the spatial distribution of traces in the parasite-host system under study. However, in our research, this approach can be applied to only 7% of the trace data, excluding the majority of the malformations (Table S2). To take advantage of the available trace data and fully explore the spatial patterns in the distribution of parasitic traces in the host species under study, it is necessary to combine malformations from multiple skeletons into spatial point patterns. Previous studies using drilling traces on bivalve prey (Rojas et al., 2015; Karapunar et al., 2023) showed that the geometric morphometric approach used in the SPPAT allows us to address that challenge. Here the assembled spatial dataset combining the locations of trematode-induced pits from multiple specimens reproduces the spatial patterns observed in a single multi-parasitized specimen. This suggests that the spatial patterns observed in the combined datasets are unlikely to be artifacts of the approach and supports their potential for evaluating the patterns in the parasite-host systems under study. In addition, we investigated the number of trematode-induced pits required to capture the dynamics of the parasite-host system under study. We found that consistent patterns emerge across a wide range of trace counts. Based on the bootstrapped spatial point patterns results, we recommend using a minimum of 25 traces to characterize the parasite-host system using SPPAT effectively. Finally, we showed how to employ point patterns of parasite-induced malformation on a shelled invertebrate host to analyze bivariate interactions among different parasite taxa in a host using distance-based statistics (e.g., CSR, aggregation, and segregation for the univariate case and independence, attraction, and repulsion in the bivariate case). Other null models can be used for testing more complex hypotheses suitable to describe other parasite-host systems, such as the area-interaction process (Badeley and Lieshout, 1995), in which points of the same type do not interact while points of different types are forbidden to lie closer than a given distance.

### 4.2. Biological implications of spatial point analysis on trematode-induced pits

We assume that the internal shell surface lacks heterogeneity, and thus, any non-homogeneity in the distribution of traces reflects the underlying interactions between point events. However, our empirical study suggests that the non-homogeneity in the distribution of traces may be attributed to specific processes controlling their spatial distribution, rather than to interactions between traces. Point patterns combining the locations of trematode-induced pits from multiple specimens replicate the spatial patterns observed in a single multi-parasitized specimen, even though the metacercariae did not interact within the same host. The spatial arrangement of the trematode-induced malformations on *C. gallina* is likely a subrogate of unobservable, and poorly understood mechanisms of the infective dynamics that regulate the specific parasite-host interactions during the life of the host. The clusters of pits arranged in a linear fashion that drive the significant segregated pattern could be indicative of such a mechanism. There is little known about the activities of trematode parasites in living *C. gallina* and the host response (see chapter 2.1). The ecological surveys of Bartoli (1974) provide some insights. *G. rostratus* metacercariae were identified in the extrapallial space between the mantle and shell wall of several bivalve second intermediate hosts including *Tapes decussatus*, *Donax truculus*, and *Solen marginatus*. Bartoli (1974) also describes *P. duboisi* as occupying the extrapallial space of their second intermediate hosts but does not provide further detail. The documented interactions of other similar parasite-host systems can also be informative. For instance, in the *Bartolius pierrei*-*Darina solenoides* gymnophallid-bivalve relationship, *B. pierrei* metacercariae are commonly embedded in a sack of mantle tissue that detaches from the mantle and migrates into the visceral mass of the bivalve, this reaction does not seem to elicit the formation of any pits. However, when the clam is heavily parasitized (i.e., the space in the visceral mass is filled), characteristic open pouches with calcifications in the form of crest or ridges represent the bivalve attempt to contain the metacercariae of *B. pierrei* and leave a characteristic trace on the interior of the bivalve exoskeleton (Cremonte 2004). Thus, the formation of such traces, in this case, represents the output of heavily parasitized aged clams where the infection process had reached more advanced stages, and its size as a taxonomical proxy is challenging to assess. In other digenean trematodes belonging to gymnophallid and other families, similar behavior is reported, but it represents the ordinary response of the host to the metacercariae, or it is associated with the metacercariae state. Campbell (1985) was one of the first authors to document that *Gymnophallus rebecqui* metacercariae living in the extrapallial space of the bivalve *Abra* (in Great Britain), which commonly induced hyperplasia of the mantle epithelium and contemporary deposition of shell material around the parasite, forming diagnostic traces having raised rims and representing the interactions of one single metacercaria (“blister”) or of a small group of them (“crater”). A somewhat similar reaction to single metacercariae was described by Ituarte et al. (2001) for a bivalve (*Gaimardia* from Argentina) in relation to an open nomenclature taxon belonging to Lepocreadiidae, a digenean trematode group that commonly has fish as the final host (Bartoli, 1974). In that case, metacercariae were recovered in the extrapallial space and lodged in a rounded or oval shallow pit. Therefore, the way species are spatially arranged, and their size classifications may not necessarily serve as a proxy for taxonomy, even at the genus level. This is contingent upon the taxonomy of the species involved in the interactions and the timing of those interactions (either initial or mature). However, gathering this information in the literature for our case study is still challenging. Our experiments, including those examining bivariate interactions, supported the theoretical null model suggesting that the two spatial point patterns are generated by independent processes, meaning that different types of size-classes do not engage with each other.

Nevertheless, the high density of traces in the dorsal half of the valve and near its ventral edge, which configures the mixed aggregated and segregated pattern, suggests a stereotypy related to the infection (i.e., host behavior and location of parasites) that agrees well with that of several taxa of digenean trematodes mainly (but not exclusively) attributed to the family Gymnophallidae from a variety of marine regions worldwide (even a more sound comparison should be made by examining traces on modern dead shells). Thus, pointing toward an identification at the large taxonomic level and, more importantly, highlighting stability in the modality of infection by means of such digenean trematode larval stages. Indeed, most of the gymnophallid (but also some Lepocreadiidae; Ituarte et al. 2001) metacercariae colonize and live within the extrapallial space which they reach after being passively ingested or actively piercing the mantle. Only for those metacercariae that, for at least a certain period of their life stage, reside in the extrapallial space, it might be possible to induce the formation of carbonate precipitation from the host. However, this phenomenon is species-specific (see the different responses of gymnophallid taxa reported by Ittuarte et al., 2001, 2009), or it could also depend on the parasite intensity (see previous section). The aggregated and marginally segregated pattern with high density in the dorsal inner area here documented is also typical of several gymnophallid genera. According to Campbell (1985), the metacercariae of *G. rebecqui* are predominantly located in the dorsal portion of the extrapallial space in both *A. tenuis* and *C. glaucum* found along the southern coast of England. This specific distribution might stem from the cercariae entering the mantle from the mantle cavity and then migrating dorsally within the extrapallial space. Alternatively, as shown with *Gymnophallus fossarum* in the brackish bivalve *C. glaucum* (see Bartoli, 1973), cercariae may penetrate the gill or labial palp epithelium, move through the tissue near the digestive gland, and then breach the mantle epithelium in the dorsal region of the bivalve. While this pathway presents greater difficulties for the cercariae, it could be crucial for any parasite that is carried along with the significant sediment consumed during feeding, transported through the ciliary tracts of the gill lamellae and labial palps. Similarly, Lepocreadiidae pit-forming metacercariae in *Gaimardia* bivalve were found in a wider area of the extrapallial space with respect to the co-occurring gymnophallid parasites that tended to prefer the inner dorsal part of the host (Ituarte et al., 2001).

### 4.3. Caveats concerning methodological assumptions underlying our modeling of pit size as a taxonomic indicator

Anecdotal evidence suggests that trematode-induced pits found in bivalves exhibit variation in both size and shape. The methodological decision to create two size classes, including the threshold of 0.55 mm, followed a previous study of the modern and late Holocene samples included in our research, indicating that both sets of pit samples are made up of two Gaussian distributions (Fitzgerald et al. 2024). Indeed, a primary factor contributing to the variability in metacercariae size is taxonomic identity. Bartoli (1974) documented the minimum and maximum lengths and widths of two trematode taxa that infect *C. gallina* near the Rhône River delta in southern France. While there is considerable overlap in body sizes at the species level, notable differences appear at the genus level, with *Parvatrema* metacercariae generally larger than *Gymnophallus*. Thus, trematode pit size may serve as a taxonomic marker, providing insight into the paleoecological connections between parasite groups and their hosts over time. Hence, pit size distributions can be utilized to investigate the richness and dynamics of specific trematode taxa, though with few underlying assumptions of this approach. First, the metacercariae growth within the bivalve second intermediate host is minimal. According to Saldanha et al. (2009), growth occurs from the moment cercariae invades the bivalve until the metacercariae fully encyst over the span of weeks, at which point growth halts (see also Cremonte, 2004). Consequently, the size of the metacercariae remains consistent even as the host grows. Second, we assume that the body size distribution of a metacercariae population follows a normal distribution. Although extensive data on this are scarce, Saldanha et al. (2009) provided size-frequency distributions for 1,510 cyst diameters of *Maritrema novazealandensis* (Microphallidae) extracted from isopods, which seem to exhibit a slight positive skew while appearing normally distributed. Third, we assume a direct relationship between trematode-induced pit size and metacercariae body size. Some studies have illustrated images of trematode metacercariae residing within the structures whose growth they induce. Ruiz and Lindberg (1989) showcased a SEM micrograph of *Parvatrema borealis* situated within a pit formed on the inner surface of *Transennella confusa* from Bodega Bay, California, where the metacercariae measured 0.149 mm (geometric mean of length and width) and occupied a significant portion of the pit’s interior width of 0.295 mm (geometric mean of primary and secondary axis). Fourth, we assume that the quantity of trematode-induced pits on a valve correlates with the abundance of metacercariae infecting a specific clam. Even if this is taxonomically driven as some groups elicit host responses for individual metacercariae (e.g., Lepocreadiidae gen. sp. Ituarte et al., 2001). For others the growth response can reflect individual or small groups of metacercariae (e.g., blister and crater-like pits in Campbell 1985). No study has examined the correlation between trematode-induced pit numbers and the number of metacercariae infecting individual in *C. gallina* hosts. Hence, the number of pits serves as a proxy for trematode abundance even if it reasonably underestimates the true amount.

## 5. Conclusions

We presented a framework for quantifying spatial patterns of trematode-induced malformations on bivalve shells using a case study on the commercially relevant bivalve *C. gallina*. Consistent with previous qualitative assessments, we found that trematode-induced malformations on bivalve shells are not random; they show an aggregated pattern for metacercaria traces of the same size classes, while an independent pattern arises when examining traces of two distinct size classes. Furthermore, our experiments show that spatial datasets combining locations of trematode-induced pits from multiple specimens reproduce the spatial patterns observed in a single multi-parasitized specimen (≥ 25 traces are recommended to characterize the parasite-host system). This suggests that patterns observed in the combined datasets are unlikely to be artifacts of the SPPAT approach and supports their potential for describing the parasite-host dynamics. This framework is applicable to a wide range of parasite-host systems in both marine and freshwater environments. It can be used to assess potential changes in the antagonistic interactions between parasites and host species over time and across space. By examining the distribution patterns of trematode-induced traces across geological time scales, researchers can investigate the long-term dynamics of parasite-host interactions (e.g., stability, stochasticity, or other proposed dynamics), providing valuable context for ecological monitoring of infection behaviors in specific parasite groups, both historically and in contemporary times. Furthermore, our approach may reveal potential environmental drivers correlated with changes in the examined biotic interactions during ecosystem alterations (e.g., Scarponi et al., 2022).

## Acknowledgments

This work was financed by the European Union–NextGenerationEU through the Italian Ministry of University and Research under Piano Nazionale di Ripresa e Resilienza (PNRR): Mission 4 Component C2, Investment 1.1 “Conservation of life on Earth: The fossil record as an unparalleled archive of ecological and evolutionary responses to past warming events.” PI Cinzia Bottini. Prot. 2022WEZR44. JWH was supported by the National Science Foundation (NSF EAR CAREER 1650745 and SGP 2409210).

## Supplementary Information

Supplementary Figure S1.

Supplementary Table S1.

Supplementary Data Captions

Supplementary R Script.

## Data availability

All data used in this research are available in the Supplementary data

