## Supplementary Information for "Spatial patterns of trematode-induced pits on bivalve skeletons: Challenges and prospects for research on parasite-host dynamics"

**This PDF file includes:**

- Supplementary Figure S1
- Supplementary Table S1
- Supplementary Legends for Supplementary Data S1 to S3.
- R script

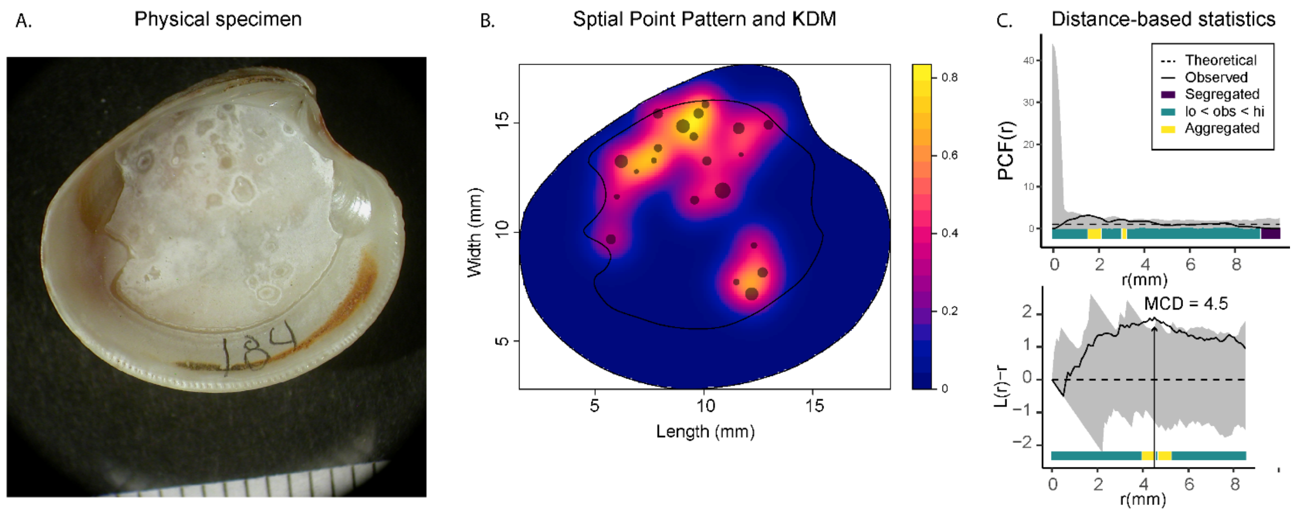


**Supplementary Figure S1.** Univariate analysis. The assembled datasets give the locations of trematode-induced pits in a single specimen. **A**. Valve 240S8 13.10m 184 L; 21 pits. **B**. Point pattern and kernel density map in standard units (millimeters). B. L-function. The black arrow indicates the point of maximum clustering distance (MCD). C. Pair correlation function (PCF) and L-function. The dark gray area is the simulation envelope for 999 Monte Carlo simulations of Complete Spatial Randomness CSR (Theoretical). The empirical curves of the distance-based statistical functions (i.e., observed data) are compared to the Monte Carlo envelopes generated through simulations of the null model. A departure from the null model is indicated when the empirical curves fall outside the simulation envelopes. The maximum clustering distance (MCD = 4.5 mm) represents the scale at which pits are most strongly clustered (i.e., the distance r at which the observed value of the L-function deviates the most from the expected value under CSR). The position of the empirical curve in relation to the simulation envelope of the null model is indicated by a color bar at the bottom. Intuitively, the kernel density map indicates an aggregated patterns with two clusters. However, given the limited reduced number of traces, the graphical outputs of the distance-based statistics show marginally significant clustering.

| # pits per valve | # valves | MCD | u* | rank | p.value | bandwidth | mc x | mc y | std x | std y |
|---|---|---|---|---|---|---|---|---|---|---|
| 1 | 62 | 0.42 | 0.0188 | 1 | 0.001 | 0.04 | 0.0541 | 0.2565 | 0.1877 | 0.1842 |
| 2 | 12 | 0.4 | 0.0137 | 1 | 0.001 | 0.05 | 0.0839 | 0.2243 | 0.1752 | 0.2314 |
| 3 | 22 | 0.41 | 0.0177 | 1 | 0.001 | 0.04 | 0.1194 | 0.2540 | 0.2112 | 0.1807 |
| 4 | 17 | 0.35 | 0.0088 | 1 | 0.001 | 0.05 | 0.0351 | 0.2134 | 0.2027 | 0.2604 |
| 5 | 14 | 0.35 | 0.0272 | 1 | 0.001 | 0.04 | 0.1277 | 0.2844 | 0.1557 | 0.1665 |
| 6 | 2 | 0.26 | 0.0178 | 7 | 0.007 | 0.06 | 0.1683 | 0.2210 | 0.1070 | 0.2555 |
| 7 | 4 | 0.4 | 0.0265 | 1 | 0.001 | 0.05 | 0.1679 | 0.2858 | 0.1468 | 0.1815 |
| 6 | 2 | 0.26 | 0.0178 | 14 | 0.014 | 0.06 | 0.1683 | 0.2210 | 0.1070 | 0.2555 |
| 8 | 4 | 0.37 | 0.0138 | 1 | 0.001 | 0.05 | 0.1305 | 0.1982 | 0.2094 | 0.1846 |
| 9 | 5 | 0.47 | 0.0142 | 1 | 0.001 | 0.05 | 0.0995 | 0.2073 | 0.1677 | 0.2330 |
| 10 | 2 | 0.33 | 0.0152 | 1 | 0.001 | 0.06 | 0.1201 | 0.1710 | 0.1852 | 0.1913 |
| 11 | 6 | 0.44 | 0.0154 | 1 | 0.001 | 0.04 | 0.1346 | 0.2246 | 0.1875 | 0.2101 |
| 13 | 2 | 0.47 | 0.0126 | 1 | 0.001 | 0.06 | 0.1666 | 0.1708 | 0.2118 | 0.1985 |
| 15 | 1 | 0.51 | 0.0236 | 1 | 0.001 | 0.06 | 0.0969 | 0.3216 | 0.1770 | 0.1812 |
| 16 | 2 | 0.38 | 0.0092 | 1 | 0.001 | 0.06 | 0.1827 | 0.1042 | 0.1462 | 0.2813 |
| 19 | 1 | 0.38 | 0.0223 | 1 | 0.001 | 0.05 | 0.1708 | 0.2449 | 0.1592 | 0.1827 |
| 21 | 1 | 0.39 | 0.0115 | 1 | 0.001 | 0.07 | -0.0329 | 0.2269 | 0.2077 | 0.2423 |
| 26 | 2 | 0.39 | 0.0119 | 1 | 0.001 | 0.05 | 0.1196 | 0.1421 | 0.1646 | 0.2269 |
| 29 | 1 | 0.36 | 0.0195 | 1 | 0.001 | 0.05 | 0.2310 | 0.2157 | 0.1778 | 0.2018 |
| 32 | 1 | 0.45 | 0.0082 | 1 | 0.001 | 0.06 | 0.1032 | 0.1558 | 0.2350 | 0.2102 |
| 34 | 1 | 0.41 | 0.0194 | 1 | 0.001 | 0.05 | 0.1487 | 0.2317 | 0.1833 | 0.1760 |
| 43 | 1 | 0.49 | 0.0097 | 1 | 0.001 | 0.05 | 0.0778 | 0.1779 | 0.2029 | 0.2307 |
| 48 | 1 | 0.35 | 0.0102 | 1 | 0.001 | 0.05 | 0.1833 | 0.1093 | 0.2000 | 0.2414 |
| 63 | 1 | 0.49 | 0.0045 | 1 | 0.001 | 0.06 | -0.0609 | 0.1081 | 0.2551 | 0.2392 |

mean center (mc); unweigted standard distance (sd); *Diggle-Cressie-Loosmore-Ford test of CSR.

**Supplementary Table S1**. Descriptive statistics of the point patterns combining data from hosts with multiple traces (i.e., multi-parasitized shells), as illustrated in Figure 8. The assembled datasets represent the locations of trematode-induced pits from all valves exhibiting the same number of traces in core 240S8. The maximum clustering distance (MCD) indicates the scale at which pits are most strongly clustered. The mean center (mc) is the average (centroid) location of all pits in the dataset. The unweighted standard distance (sd) measures the average dispersion of points around the mean center, reflecting how spread out the points are and summarizing the spatial variability of the distribution.

Supplementary Data captions

**Supplementary Data S1.** Assembled point data on trematode-induced pits on *C. gallina*. This dataset provides the locations and sizes of trematode-induced pits in the examined valves. It includes the original landmark coordinates as well as the rotated, scaled, and translated coordinates obtained through Bookstein baseline registration using landmarks 3 and 6.

**Supplementary Data S2.** Outer area. This dataset provides rotated, scaled, and translated coordinates of the observation window obtained through Bookstein baseline registration using landmarks 3 and 6.

**Supplementary Data S3.** Internal area. This dataset provides rotated, scaled, and translated coordinates of the area used as a reference, obtained through Bookstein baseline registration using landmarks 3 and 6.
